# Published Anti-SARS-CoV-2 In Vitro Hits Share Common Mechanisms of Action that Synergize with Antivirals

**DOI:** 10.1101/2021.03.04.433931

**Authors:** Jing Xing, Shreya Paithankar, Ke Liu, Katie Uhl, Xiaopeng Li, Meehyun Ko, Seungtaek Kim, Jeremy Haskins, Bin Chen

**Affiliations:** Department of Pediatrics and Human Development, Michigan State University, Grand Rapids, Michigan, USA; Zoonotic Virus Laboratory, Institut Pasteur Korea, Seongnam, South Korea; Department of Pharmacology and Toxicology, Michigan State University, Grand Rapids, Michigan, USA

## Abstract

The global efforts in the past few months have led to the discovery of around 200 drug repurposing candidates for COVID-19. Although most of them only exhibited moderate anti-SARS-CoV-2 activity, gaining more insights into their mechanisms of action could facilitate a better understanding of infection and the development of therapeutics. Leveraging large-scale drug-induced gene expression profiles, we found 36% of the active compounds regulate genes related to cholesterol homeostasis and microtubule cytoskeleton organization. The expression change upon drug treatment was further experimentally confirmed in human lung primary small airway. Following bioinformatics analysis on COVID-19 patient data revealed that these genes are associated with COVID-19 patient severity. The expression level of these genes also has predicted power on anti-SARS-CoV-2 efficacy in vitro, which led to the discovery of monensin as an inhibitor of SARS-CoV-2 replication in Vero-E6 cells. The final survey of recent drug-combination data indicated that drugs co-targeting cholesterol homeostasis and microtubule cytoskeleton organization processes more likely present a synergistic effect with antivirals. Therefore, potential therapeutics should be centered around combinations of targeting these processes and viral proteins.

## Main Text

As of February 15th, 2021, SARS-CoV-2 has infected 108 million people and claimed 2 million lives. Vaccines are promising for a cure, yet the emerging mutations of SARS-COV-2 impose challenges to target the virus proteins; thus targeting host cells remains a viable therapeutic approach. The global efforts in the trailing months have led to the discovery of at least 184 drug repurposing candidates *in vitro* (Supplementary Table 1). Although the majority of them only exhibited moderate antiviral potential, gaining additional insights into their mechanisms will facilitate an enhanced understanding of infection and the development of better therapeutics for COVID-19. Through integrative bioinformatics analysis of transcriptomic profiles and *in vitro* experiments, we find those positive compounds regulate cholesterol homeostasis and microtubule cytoskeleton organization pathways, which also align with COVID-19 patients severity, compound efficacy and synergism with antiviral drugs (Figure 1A).

**Figure 1.**
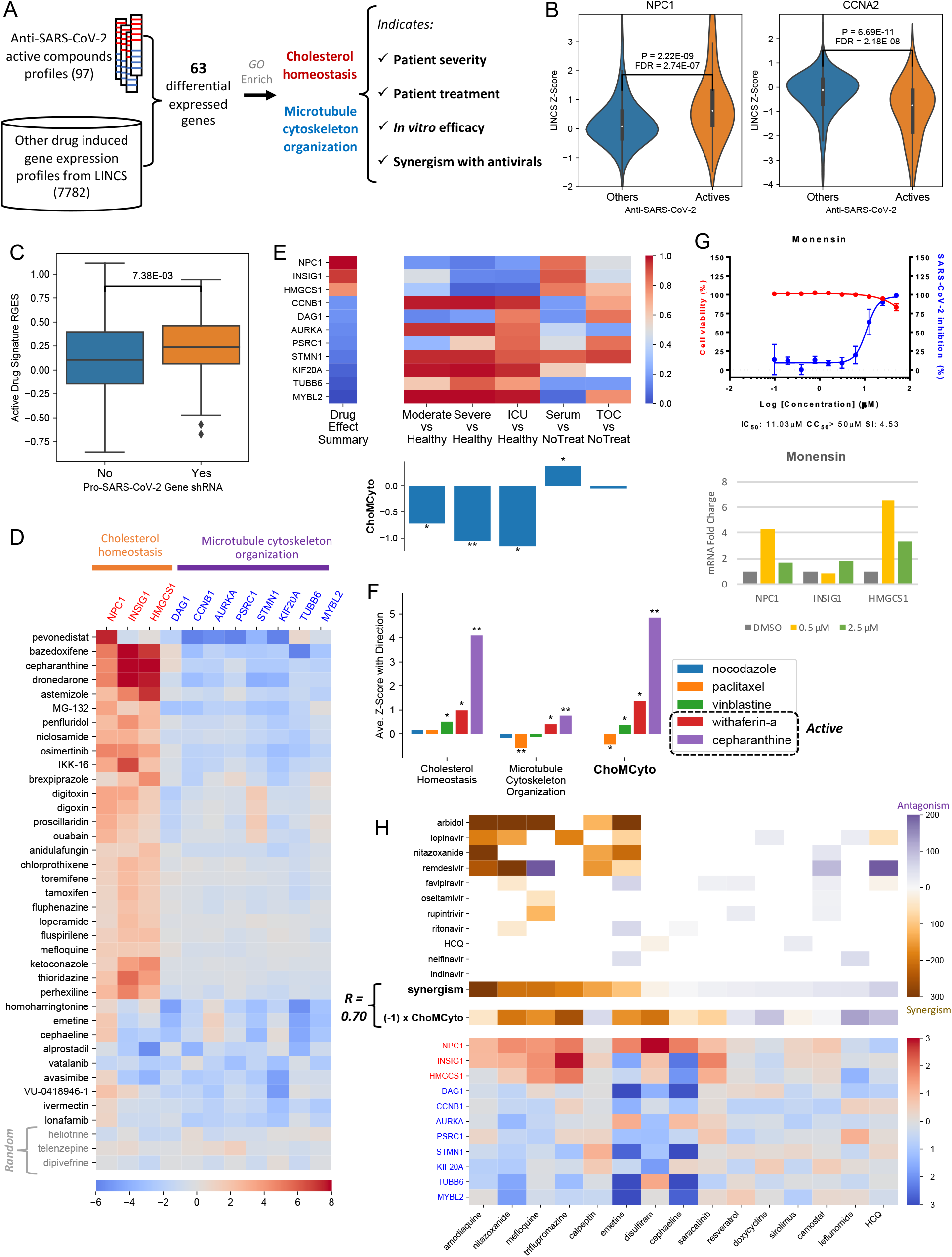
Reported anti-SARS-CoV-2 compounds regulate genes related to cholesterol homeostasis and microtubule cytoskeleton organization, which are associated with COVID-19 patients’ severity and synergism with antivirals. **A**, the workflow of this research. **B**, Example genes were induced or suppressed by anti-SARS-CoV-2 compounds. The Y-axis indicates the LINCS z-score of a specific compound and a higher score means higher expression change. P values were derived from Wilcox rank-sum tests, and further corrected across all LINCS 978 genes. **C**, A boxplot comparison between anti-SARS-CoV-2 CRISPR screening gene hits and non-hits. A higher RGES score on Y-axis indicates a query host cell shRNA knockdown-induced gene expression profile more closely resembles the summarized gene expression signature of anti-SARS-CoV-2 active compounds. X-axis denotes whether a query host gene knock-out makes the cells resistant to SARS-CoV-2 infection. **D**, Expression of genes involved in cholesterol homeostasis or microtubule cytoskeleton organization was changed by 38 anti-SARS-CoV-2 compounds but not by three randomly selected compounds namely heliotrine, telenzepine and dipivefrine. **E**, The expression change of the selected ChoMCyto genes in different COVID-19 patient groups. In the heatmap, the log2 fold change values from different comparisons were converted into gene rankings within a transcriptome. The overall ChoMCyto scores are shown in the bar plot. TOC: tocilizumab. Patient severity dataset: SRP267176; treatment dataset: SRP301622. **F**, Drug effect on ChoMCyto genes is associated with the antiviral efficacy *in vitro*. **G**, Top: the dose-response (blue) and dose- viability (red) curves of monensin, with IC_50_, CC_50_, and selectivity index labeled. Bottom: mRNA fold changes (compared with TBP) of NPC1, INSIG1 and HMGCS1 induced by DMSO (control), 0.5 µM monensin or 2.5 µM monensin. **H**, The drug effect on ChoMCyto genes correlates with antiviral synergism. The top and bottom panels share drug columns. HCQ: hydroxychloroquine. The top panel illustrates synergistic effects between host-targeting drugs and antivirals (white: missing values, purple: synergism, orange: antagonism). The last row “Synergism” summarizes the average synergistic effect of antiviral drugs in all rows for each specific compound in the X-axis. The middle row illustrates the ChoMCyto scores multiplied by -1, for a better color agreement with the row above it. The bottom panel shows the expression change of ChoMCyto genes induced by each drug. In all the heatmaps except the synergistic one, red indicates up-regulation, and blue means down-regulation. Gene symbols colored with red are up- regulated by anti-SARS-CoV-2 compounds, and those colored with blue are down-regulated. *: p < 0.05; **: p < 0.001. P values were derived from randomly shuffled signatures permutation (Supplementary Methods).

Leveraging large-scale drug-induced gene expression profiles from the Library of Integrated Network-Based Cellular Signatures (LINCS) project ^1^, we computed the differential gene expression induced by 97 anti-SARS-CoV-2 positive compounds versus 7,782 others, which identified 63 genes specifically dysregulated (30 down-regulated and 33 up-regulated, Supplementary Methods, Supplementary Table 2). For example, *NPC1*, an intracellular cholesterol transporter, is highly up-regulated (z score >= 2) by 17 active compounds, much higher expression than in other compounds (p = 2.2E-09, Wilcox rank sums test, Figure 1B). In addition, the expression change of *NPC1* is positively correlated with drug EC_50_ (Spearman Correlation Rho = -0.44, p = 8.2E-05, Supplementary Table 2). The expression change upon drug treatment was further confirmed in human lung primary small airway cells (Supplementary Figure S1A). An opposite pattern exists in *CCNA2*, a G1/S and G2/M transition regulator (p = 6.69E-11, Rho = 0.50, Figure 1B and Supplementary Table 2).

The 63 genes differential expression might serve as a cellular response signature for anti-SARS-CoV-2 candidates. We then validated this signature with five independent genome-wide CRISPR screening datasets from ^2–6^. These studies identified 49 host factor genes critical in SARS-CoV2- infection (Supplementary Table 3), termed pro-viral genes. Assuming that knock-down of individual pro-viral genes might benefit the host cell in SARS-CoV-2 challenging, we computed the similarity between the anti-SARS-CoV-2 compound signature and the expression profiles perturbed by shRNA of each pro-viral gene through a gene-set enrichment analysis (adopted from RGES) ^7^. The RGES values of 49 pro-virus genes knock-down are significantly higher than the remaining 4321 genes (p = 7.38E-03, Wilcox rank sums test, Figure 1C), suggesting inhibition of pro-viral genes has similar effects with the active compounds, and the anti-SARS-CoV-2 signature captures key biological processes involved in viral infection.

By incorporating the gene co-expression knowledge, we performed Gene ontology (GO) enrichment analysis with a background beyond LINCS 978 genes. For the 33 up-regulated genes, cholesterol biosynthetic and metabolic processes (“cholesterol homeostasis” ^8^ hereafter) were enriched, with genes *NPC1, INSIG1* and *HMGCS1* involved (Supplementary Figure S2A). For the 30 down-regulated genes, the “microtubule cytoskeleton organization” process was enriched, with genes *DAG1, CCNB1, AURKA, PSRC1, STMN1, KIF20A, TUBB6*, and *MYBL2* annotated to this term (Supplementary Figure S2B). Although mitotic cell cycle related pathways were significant, we didn’t further investigate them because of the biased distribution of the cancer enriched genes in the LINCS data. Overall, 36% of the active compounds change the expression of the cholesterol homeostasis or microtubule cytoskeleton organization pathway members (average z score >= 1.5, Figure 1D). Strikingly, these compounds are not antivirals, and their primary mechanism of action varies, such as NF-kB inhibitors, selective estrogen receptor modulators and histamine receptor antagonists. Therefore, their antiviral activity might be an off-target or indirect effect on cholesterol homeostasis and/or microtubule cytoskeleton organization.

Next we examined the expression of the genes involved in the two pathways using two independent COVID-19 patient cohorts (33 samples from PBMC and 50 samples from blood). As shown in Figure 1E and Supplementary Table 4, the comparison between the patient and healthy group showed a reversal pattern to the summarized active drug effect, while the transcriptional changing pattern of convalescent serum treatment was in line with active compounds. We designed a <u>C</u>holesterol <u>ho</u>meostasis and <u>M</u>icrotubule <u>Cyt</u>oskeleton <u>o</u>rganization (ChoMCyto) score to quantify the reversal (negative) or mimicking (positive) pattern. This score associates with patients’ severity and treatment. Of note, tocilizumab treatment did not affect these genes (ChoMCyto approximately zero), probably because it’s an immune-suppressor, not aiming to target viral infection processes. Together, cholesterol homeostasis and microtubule cytoskeleton organization pathways are associated with COVID-19 severity, and co-targeting these two pathways may improve the outcome.

We then investigated if the ChoMCyto score has predictive power on anti-SARS-CoV-2 activity. Cortese et al. ^9^ found SARS-CoV-2 caused cytoskeleton remodeling imperative for viral replication. After testing a few drugs altering cytoskeleton integrity and dynamics, they only observed withafterin A had a robust antiviral effect. Gene expression profiles of these drugs suggested they acted differently on ChoMCyto genes, although targeting cytoskeleton proteins. Withaferin A showed a similar pattern with the active drugs, while paclitaxel showed a significantly opposite pattern (Figure 1F). In addition, withaferin A strongly regulated cholesterol homeostasis, while the other three didn’t (Figure 1F); thus, withaferin A achieved a higher ChoMCyto score. A stronger pattern was observed for the positive control, cepharanthine (EC_50_ = 4.47 µM ^10^). It was reported blocking cholesterol trafficking via targeting NPC1 ^11^. We also observed that cepharanthine inhibited actin expression in human lung primary small airway cells (Supplementary Figure S1B). Further, ChoMCyto score was applied to all LINCS compounds (Supplementary Table 5). Among the FDA-approved drugs, two top candidates, lomitapide (MTTP inhibitor for hypercholesterolemia treatment) and monensin (an ionophore reported anti-MERS activity) were not evaluated yet, thus selected to test anti-SARS-CoV-2 activity *in vitro*. We found that monensin inhibited SARS-CoV-2 replication in Vero cells with IC_50_ of 11 µM, and its CC_50_ was > 50 µM (Figure 1G). It also induced expression of *NPC1, INSIG1* and *HMGCS1* in small lung airway cells (Figure 1G). Although lomitapide was inactive under this experimental setting (Supplementary Figure S3), Mirabelli et. al.^12^ reported its IC_50_ as 765 nM in Huh7 cells. In addition, among the top 20 candidates, bazedoxifene, dronedarone and osimertinib were already reported active ^13^. This suggests that regulating cholesterol homeostasis and microtubule cytoskeleton organization might contribute to antiviral efficacy. Comparison between cytotoxic (CC_50_ < 50 µM) and non-toxic hits suggests that ChoMCyto genes expression change doesn’t significantly contribute to the cytotoxicity (Supplementary Table 6).

Since SARS-CoV-2 entry and infection of cells comprises multiple critical biological processes inside of infected cells, we further evaluated the combination of targeting ChoMCyto genes and other processes such as viral replication in COVID-19 treatment. To do so, we elicited a recent combination screening study from NCATS ^14,15^, where 15 host-targeting compounds were combined with at least one of 11 antivirals (9 known antivirals, two potent SARS-CoV-2 candidates nitazoxanide and hydroxychloroquine) (Figure 1H, Supplementary Table 7). For each host-targeting compound, we summarized its synergistic effect with the 11 antivirals (see Methods) as well as the ChoMCyto score. The drugs with higher ChoMCyto scores are more likely to present a synergistic effect with antivirals (Spearman correlation of -0.70, Supplementary Figure S4). For instance, the melfloquine effect on ChoMCyto genes shares a similar pattern with reported anti-SARS-CoV-2 active compounds, and it strongly synergizes with arbidol, a viral envelope fusion inhibitor. While leflunomide has a negative ChoMCyto score, and antagonizes with lopinavir or nelfinavir. Remdesivir showed a synergistic effect with amodiaquine, nitazoxanide and emetine, it may retain a potential synergistic effect with triflupromazine, which was not tested yet, but with the highest ChoMCyto score. Among the top of all repurposing candidates, cepharanthine (ChoMCyto = 9.44, Supplementary Table 5) was also reported synergism with remdesivir, that 2.5 µM of both could inhibit > 95% cytopathic effect *in vitro*, while only 35% and 10% for each single agent ^14^. Although either agent alone only presented weak antiviral activity (EC_50_ at micromolar level), their combination exerted a marked effect. This suggests the therapeutic potential of the anti-inflammatory drug cepharanthine combined with antiviral treatment such as remdesivir.

In summary, our survey of the positive hits from screenings reveals they share common mechanistic effects through the regulation of cholesterol homeostasis and microtubule cytoskeleton organization. We designed a ChoMCyto score to quantify this summarized drug effect pattern, which is associated with COVID-19 patient severity and treatment. By applying the ChoMCyto pattern to predict anti-SARS-CoV-2 efficacy, we discovered monensin with EC_50_ of 11 µM *in vitro*. Literature survey suggested the antiviral mechanism of monensin might be via blocking viral transport within the Golgi complex ^16^. This indirect antiviral mechanism inspired us to investigate the synergism between established antiviral drugs and repurposed drugs targeting host cellular ChoMCyto genes. Our findings corroborate the emerging evidence from other studies. For instance, the analysis of clinical data reports hypolipidemia is associated with the severity of COVID-19 ^17^. Zang et. al. ^18^ found that cholesterol 25-hydroxylase suppresses SARS-CoV-2 replication by blocking membrane fusion. In addition, its product 25-hydroxycholesterol was elevated in a fatal COVID-19 patient and infected mice ^19^. The large-scale profiling of SARS-CoV- 2 virus-host interactions found that viral proteins Orf8 and Orf9c directly bind to NPC2 and SCAP ^20^, important components for cholesterol transport and monitoring. In addition, host protein targets of viral NSP10 and NSP13 are enriched with microtubule-based process ^20^. In our recent work ^21^, 10 out of 11 ChoMCyto genes showed a contrasting expression pattern with that dysregulated by coronavirus infection. Together, we conclude that cholesterol homeostasis and microtubule cytoskeleton organization pathways are disrupted by viral infection, and anti-SARS-CoV-2 compounds tend to restore these biological processes inside the host cells. Co-targeting the two pathways might boost the efficacy of known antiviral drugs for COVID-19 treatment.

## Acknowledgments

We thank Drs. Jeremy W Prokop, Caleb P Bupp and Surender Rajasekaran for providing patient information of this dataset (SRP301622). The research is supported by R01GM134307, K01 ES028047, Spectrum Health-MSU Alliance Corporation and the MSU Global Impact Initiative. The content is solely the responsibility of the authors and does not necessarily represent the official views of sponsors.

