## Supplementary Materials for "Published Anti-SARS-CoV-2 In Vitro Hits Share Common Mechanisms of Action that Synergize with Antivirals"

### **Figures**

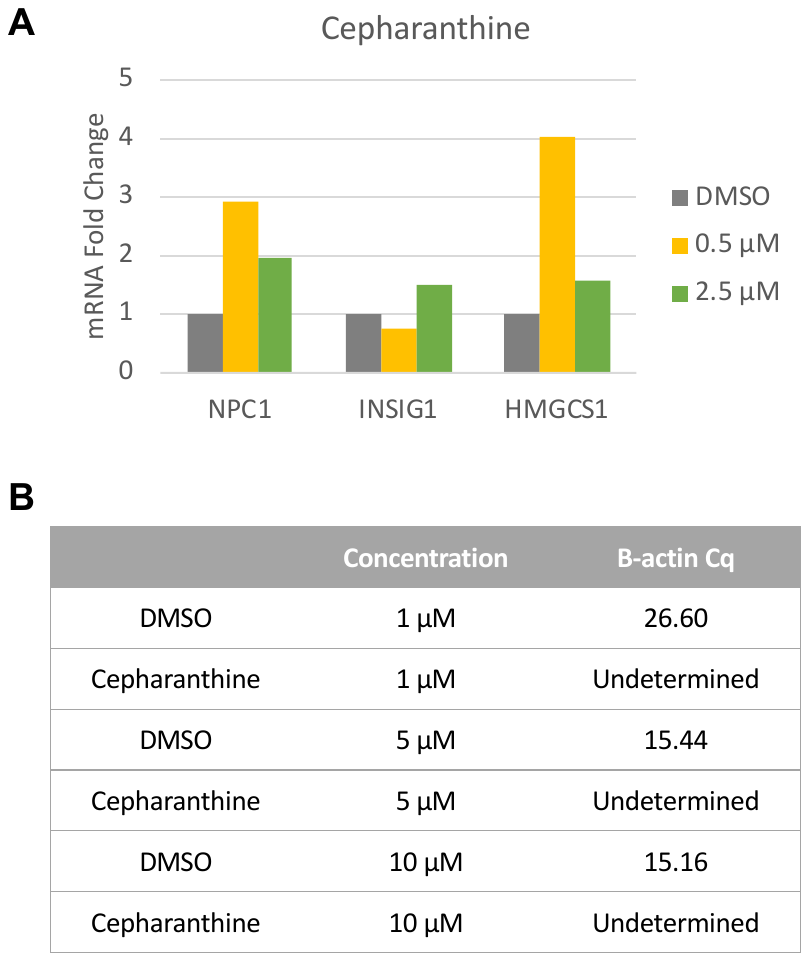

Figure S1. Gene expression change induced by cepharanthine in human lung primary small airway cells. A, The expression change of NPC1, INSIG1 and HMGCS1. The Y axis indicates the amount of mRNA fold change for a specific gene (e.g. NPC1) compared with TBP. B, Beta-actin expression was inhibited by cepharanthine treatment. Cq, quantification cycle.

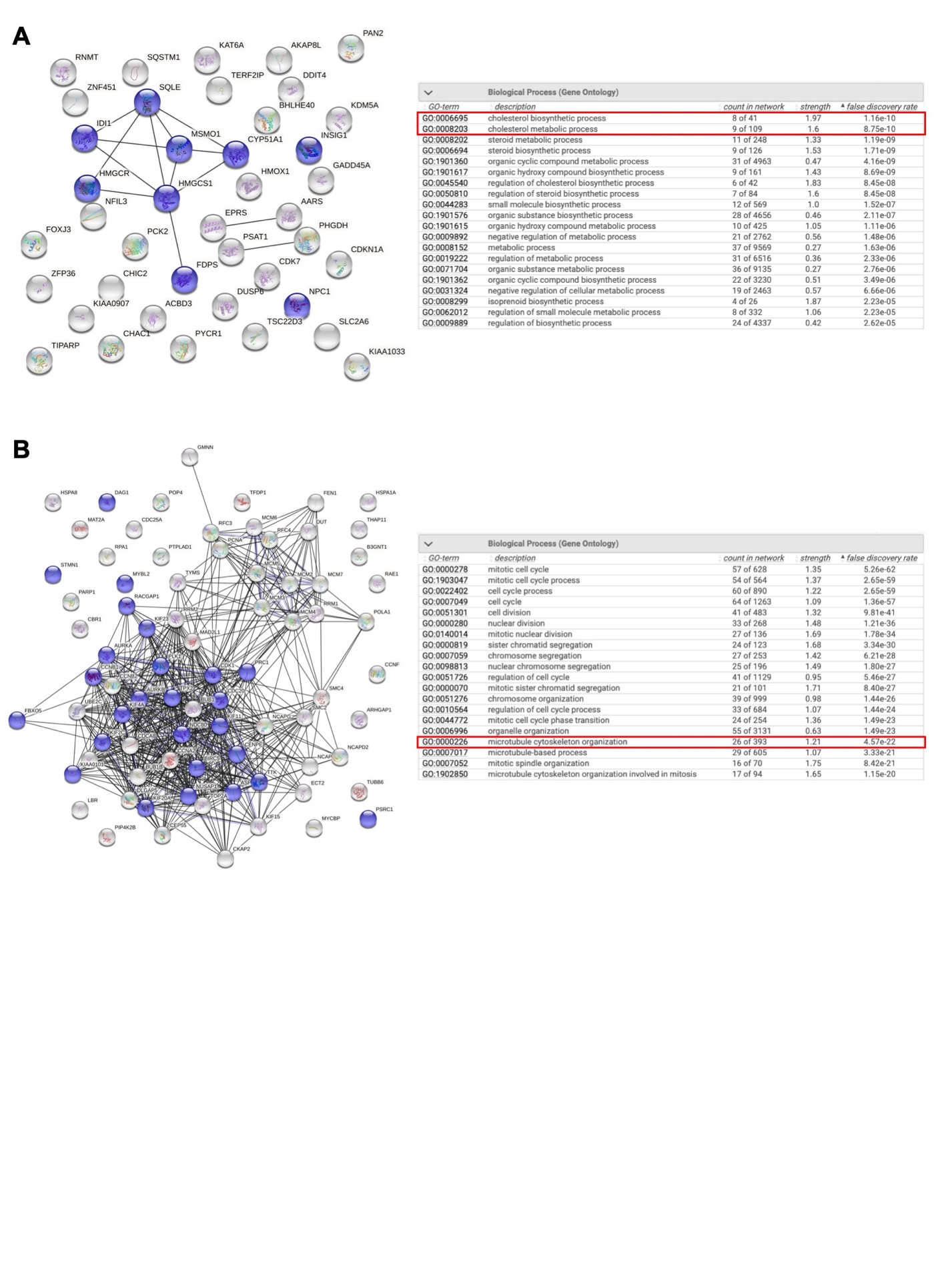

Figure S2. Gene Ontology (GO) enrichment results of up-regulated (A) and down-regulated (B) genes. Pathways of interest are highlighted. Their members in the networks are colored in purple.

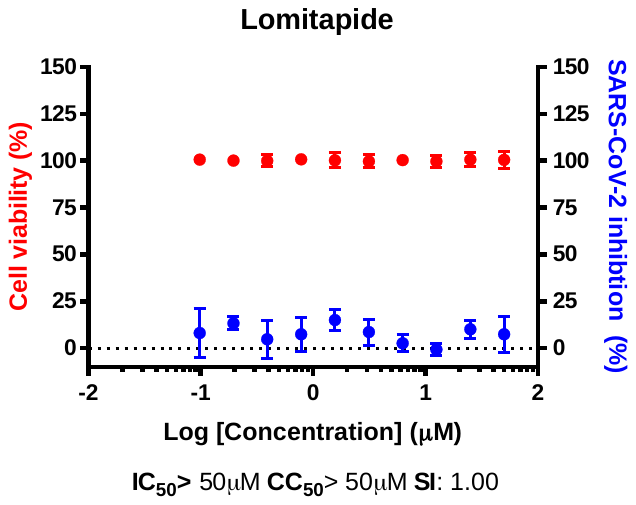

Figure S3. Dose-response curves of lomitapide. Red points indicate cell viability changing with different dose applied. Blue points indicate SARS-CoV-2 inhibition.

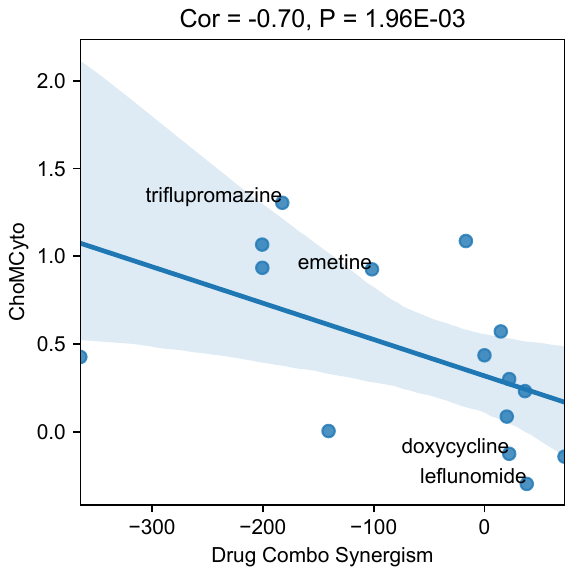

Figure S4. ChoMCyto score is correlated with compound-antivirals average synergistic effect.

### **Methods**

#### **Average compound or shRNA induced gene expression changes**

The drug-induced gene expression profiles produced in the LINCS L1000 project^1^ are accessible in GEO with IDs GSE92742 (Phase 1) and GSE70138 (Phase 2). The level-5 profile derived from the comparison of gene expressions between the perturbagen- or vehicle control-treated samples represents gene expression changes upon compound or shRNA treatment. Only 978 landmark genes and high-quality profiles (is_gold = 1, annotated in the meta information) were included in the analysis. As one compound could be profiled under different concentrations, treatment durations, and cellular contexts, for a specific compound, we took the median LINCS z-scores of its profiles measured at 10 µM of drug treatment, regardless of the time and cellular context, resulting in a matrix of 978 genes by 7879 compounds at the end. Similarly, we took the median z-scores of each shRNA across different testing conditions. Of note, only “CGS” shRNA profiles were used unless no “CGS” profiles were measured for a specific shRNA perturbagen. This resulted in 4370 different shRNA profiles after merging. A z-score larger than 1.5 indicates that the drug can up-regulate the gene expression, while a z-score less than -1.5 means down-regulation. A larger absolute z-score means a higher magnitude of expression change. The LINCS profiles have been extensively explored for therapeutic discovery in our previous works^2–5^.

#### **Evaluate genes expression affected by anti-SARS-CoV-2 active compounds**

The compounds profiled in LINCS were divided into two groups: anti-SARS-CoV-2 actives and others. The in vitro anti-SARS-CoV-2 efficacy data (Supplementary Table 1) were collected from the literature and the NCATS OpenData Portal (<https://opendata.ncats.nih.gov/covid19/assay?aid=14>, accessed by Jul. 2^nd^, 2020). Given the concern about data quality from high-throughput screenings, for the efficacy data from NCATS, only compounds with AC_50_ less than 5 µM were included as actives. For each landmark gene, its expression z-scores were compared between actives and other compounds using both the Wilcox rank sums test and Fisher exact test. For the Fisher exact test, the following 2 by 2 table was calculated to derive the p values of up-/down-regulation, respectively. The smaller p-value between up- and down-regulation was selected as the final p-value for this gene.

| Number of active compounds up-/down- regulating a gene | Total number of actives |
| --- | --- |
| Number of other compounds up-/down- regulating a gene | Total number of compounds not annotated as an active (other compounds) |

Then we used the Benjamini–Hochberg false discovery rate (FDR) to correct p values derived from Wilcox rank sums and Fisher exact test, respectively. The genes with both p values less than 0.05 and both FDRs less than 0.05 were chosen as significant genes affected by anti-SARS-CoV-2 drugs.

**Interpret the active compounds induced gene expression signature**

To gain the biological insights of the significant genes commonly regulated by active anti-SARS-CoV-2 compounds, we investigated their Gene Ontology (GO) enrichment with STRING-db (<https://string-db.org/>). Enrichment analysis for up-regulated or down-regulated were performed separately. As the LINCS profiles only measured 978 landmark genes expression change, which is only a small part of the whole transcriptome, we incorporated the gene co-expression knowledge to overcome this limitation. For each part of the dysregulated gene set, a co-expression network was retrieved with the highest confidence interaction score from STRING-db, with no more than 50 interactors of the second shell included. Then the GO enrichment analysis was performed on the interaction network with the whole transcriptome as background.

**Validate active compounds signature with CRISPR screening datasets**

We first compiled a pro-viral genes list based on five recent studies on CRISPR screening for anti-SARS-CoV-2 host factor genes ^6–10^. Based on the description from these papers, the following criteria were applied to define the hits. Daniloski et. al.: FDR < 0.1 and lfc (log2 fold change) > 1 for either MOI 1 or MOI 3; Hoffmann et. al.: FDR < 0.1 with a positive z-score; Schneider et. al.: FDR <0.05 and z-score > 4; Wang et. al.: score < 10E-04; Wei et. al.: Cas9-v2 average > 3. These hits were mapped to the LINCS shRNA collection. For each LINCS shRNA perturbagen, its average profile was summarized and calculated RGES to evaluate whether it could mimic (a positive RGES score) the anti-SARS-CoV-2 active compound induced gene expression change. Finally the Wilcox rank sums test was performed to compare the difference in RGES scores between the knock-down of pro-viral genes and other genes.

**Evaluate genes expression affected by anti-SARS-CoV-2 active compounds in human lung primary small airway cells**

Human small airway epithelia at the air-liquid-interface (ALI) was cultured using the same protocol for pig small airway culture as published before^11^. Human donor lungs were collected from Spectrum Health Accelerator of Research Excellence (SHARE) protocol with IRB approval. All human samples were de-identified and without personal health identifiers.

Briefly, small airway tissue is dissected from distal lungs from within 2cm from the edge of lung parenchyma. Small airways tissue was digested with pronase and epithelial cells were co-cultured with irradiated feeder cells. Expanded small airways cells were seeded on transwells for at least 2 weeks before it is ready for usage. Anti-SARS-CoV-2 active compounds were added to small airways cells with different doses for 24 hours, DMSO was used as vehicle control. RNA was isolated based on the TRIzol™ Reagent (Invitrogen, 15596026) protocol. To the 1 ml of Trizol homogenized samle, 200 µL of chloroform were added and thoroughly mixed by shaking before incubating on ice for 2-3 minutes. Samples were centrifuged for 15 minutes at 12,000 x g at 4°C. The aqueous phase was removed and transferred to a new sample tube. One volume of 70% ethanol was added to each sample and the entire contents was used for the RNA extraction process. RNA was extracted using the PureLink™ RNA Mini Kit (Invitrogen, 12183020) and eluted in 30 µL of RNase-free water. cDNA from the extracted RNA was synthesized using the SuperScript™ IV VILO™ Master Mix (Invitrogen, 11756050) with the recommended ezDNase enzyme treatment. The resulting cDNA was quantified and diluted to a concentration of 1 µg/µL. A master mix was prepared for each set of primers by utilizing the TB Green® Premix Ex Taq II (TLi RNaseH Plus) reagent kit (Takara Bio, RR820A). Primers were designed and ordered through Sigma-Aldrich (Table S1). To prepare the reaction master mix, the recommended protocol for the StepOnePlus Real-Time PCR System in the product manual was followed. Briefly, the TB Green® Premix Ex Taq II (TLi RNaseH Plus) was diluted to a final concentration of 1X and the final concentration for each primer was 0.4 µM. Except for the template, 18 µL of the master mix was added to the appropriate well of a 0.2mL Non-skirted Black-Lettered 96-well PCR plate (Thermo Scientific, AB-0600-L). Of the prepared 1 µg/µL template cDNA, 2 µL was added to each well in the plate and mixed gently. The plate was covered and centrifuged briefly to remove air bubbles from the wells. The reaction was run using a QuantStudio™ 3 Real-Time PCR System (Applied Biosystems™, A28567). Raw data was uploaded to the Thermo Fisher Cloud Dashboard for data analysis. Gene expression relative to the TBP housekeeping gene was preformed using the ΔΔCT method.

Table S1: RT-PCR Primers and Sequences

| **Gene Name** | **Sequence (5’-3’)** |
| --- | --- |
| Human NPC1 Forward | GCAGTGCCTACCGAGTATTT |
| Human NPC1 Reverse | GACACACCGAGGTTGAAGATAG |
| Human INSIG1 Forward | TCGGGCTGACTGTACAAATG |
| Human INSIG1 Reverse | CAAGGGAAGACTTCAGGGTAAG |
| Human HMGCS1 Forward | AGCCACTTTCTGTCCTCTTG |
| Human HMGCS1 Reverse | CCTGTTACCCTTGCTCTGTT |
| Human ACTB Forward | GGATCAGCAAGCAGGAGTATG |
| Human ACTB Reverse | AGAAAGGGTGTAACGCAACTAA |
| Human TBP Forward | TGTGCACAGGAGCCAAGAGT |
| Human TBP Reverse | ATTTTCTTGCTGCCAGTCTGG |

#### **Process patient blood transcriptome samples and compare between patient groups**

Raw FASTQ files for SRP301622 (https://trace.ncbi.nlm.nih.gov/Traces/sra/?study= SRP301622) and SRP267176 (https://trace.ncbi.nlm.nih.gov/Traces/sra/?study=SRP267176) were downloaded from public database NCBI SRA (https://www.ncbi.nlm.nih.gov/sra). Data were processed using RSEM^12,13^ 1.3.1 + STAR^14^ 2.6.1 pipeline. Log2 transformed (addition of pseudocount 1) TPM values as gene expression measures were used for analysis. The RNA-Seq processing code is available at GitHub (<https://github.com/Bin-Chen-Lab/chenlab_toil>). Sample meta data were obtained from GEO (GSE152418) and https://www.ncbi.nlm.nih.gov/Traces/study1/?acc=SRP301622&go=go. Gene expression differences were compared between healthy donors (control samples) and moderate/severe/ICU patients (case samples) using dataset SRP267176, and comparison between non-treated (control samples) and convalescent-serum/tocilizumab (case samples) was derived from dataset SRP301622., Lethal samples were excluded because the multi-organ dysfunction could bring dramatically change beyond viral infection. The log2 fold change values were calculated with the diffExp function in the OCTAD ^3^ R package.

**Quantify the alignment between a gene set expression pattern and a disease/drug signature**

Given expression changing pattern of a set of genes, i.e. significantly dysregulated genes together with their directions of changing (up/down-regulated), such as the ChoMCyto pattern, we calculated how strongly it matched with a disease signature (e.g. moderate patients vs. healthy donors) or a drug profile (e.g. the averaged LINCS signature of withaferin-A). In detail, es_up was derived by averaging the log2 fold change or LINCS z-score values of the genes supposed to be up-regulated in our gene set of interest. Similarly, es_down was derived by averaging the down-regulated genes of interest. Then the final score was defined as es_up – es_down. A positive score indicates the disease signature or drug profile exhibits similar pattern with the gene set of our interest (e.g. ChoMCyto genes direction summarized from active anti-SARS-CoV-2 compounds), while a negative score indicates the reversal of this pattern. A p value was derived by scoring 5000 randomly shuffled signatures.

**Evaluate inhibition of SARS-CoV-2 replication *in vitro***

We evaluated whether the proposed compounds could inhibit SARS-CoV-2 replication in an immunofluorescence assay as previously reported ^15^. In brief, Vero cells were seeded at 1.2 × 10^4^ cells per well in the 384-well-plate 24h prior to infection. Cells were treated with different concentrations (0.1 to 50 µM, 10 points) of compounds approximately 30min prior to infection. Then the cells were infected with SARS-CoV-2 (strain: βCoV/Korea/KCDC03/2020, NCCP43326) at a multiplicity of infection (MOI) of 0.01, followed with incubation at 37 °C for 24h. After fixation at room temperature for 30min and permeabilization with 0.25% tritonX-100 for 10min, the primary antibody, anti-SARS-CoV-2 Nucleoprotein (Sino Biological, 40143-T62), was attached at 37 °C for 1.5h. Then the secondary antibody attachment using goat-anti-rabbit-IgG-alexa-488 + Hoechst 33342, and nucleus staining were conducted at 37 °C for 1h. Then the assay plate was submitted to the Operetta microscope (Perkin Elmer) for imaging. The Columbus software (Perkin Elmer) was used to calculated the infectivity and cell number. Cell viability was measured by comparing the cell number to mock infection. Finally, the dose-response curve was generated with Prism 7 (GraphPad) to calculate IC_50_ with the four-parametric nonlinear fitting algorithm.

#### **Associate drug combination synergism and drug-induced gene expression**

Bobrowski et. al.^16^ measured the anti-SARS-CoV-2 synergism of 73 drug combinations in vitro. As the calculated synergistic (HSA. Neg) and antagonistic (HSA. Pos) effect separately, we took the one with larger absolute value as the synergism score of each drug combination. We investigated those pairs comprising one known antiviral drug (e.g. remdesivir and arbidol) and one non-typical-antiviral drug (e.g. amodiaquine and mefloquine). For each of the latter drugs, its general synergistic ability was summarized as an average synergism value across all the known antiviral drugs tested. Accordingly, the ChoMCyto score of these non-typical-antiviral drugs were calculated based on their LINCS profiles. Then we calculated the Spearman correlation between their average synergism values and ChoMCyto scores.

**Compare genes expression change induced by cytotoxic and non-toxic compounds**

The positive anti-SARS-CoV-2 hits from screenings were annotated as cytotoxic if reported CC_50_ less than 50 µM. For each landmark gene, its expression z-scores were compared between cytotoxic and non-toxic compounds using the Wilcox rank sums test. Then we used the Benjamini–Hochberg false discovery rate (FDR) to correct p values.

#### **Software tools and statistical methods**

All analyses were conducted in Python v3.7.6 programming language. Data matrix merge and basic statistics were performed with the Pandas package (v1.0.1). LINCS compound profile extraction was conducted using the cmapPy package (v4.0.1). Wilcox rank sums test, Fisher exact test, and Spearman correlation were calculated with the Scipy package (v1.4.1). FDR values were calculated with the Statsmodels package (v0.11.0). Data visualization was implemented with the Matplotlib (v3.1.3) and Seaborn (v0.10.0) package.
